# A single oral immunization with a replication-competent adenovirus-vectored vaccine protects mice from influenza respiratory infection

**DOI:** 10.1101/2021.07.21.453241

**Authors:** Emeline Goffin, Silvio Hemmi, Bénédicte Machiels, Laurent Gillet

**Author notes:** Address correspondence to Laurent Gillet.

## Abstract

The development of effective and flexible vaccine platforms is a major public health challenge as recently highlighted by the COVID-19 pandemic. Adenoviruses (AdVs) are easy to produce and have a good safety and efficacy profile when administered orally as demonstrated by the long-term use of oral AdV 4 and 7 vaccines in the US military. These viruses therefore appear to be the ideal backbone for the development of oral replicative vector vaccines. However, research on these vaccines is limited by the ineffective replication of human AdVs in laboratory animals. The use of mouse AdV type 1 (MAV-1) in its natural host allows infection to be studied under replicative conditions. Here, we orally vaccinated mice with MAV-1 vectors expressing the full length or the “headless” hemagglutinin (HA) of influenza to assess the protection conferred against an intranasal challenge of influenza. We showed that while the headless HA vector did not generate a significant humoral or cellular immune response to influenza, a single oral immunisation with the full-length HA vaccine generated influenza-specific and neutralizing antibodies and completely protected the mice against clinical signs and viral replication.

**Importance:** Given the constant threat of pandemics and the need for annual vaccination against influenza and possibly emerging agents such as SARS-CoV-2, new types of vaccines that are easier to produce and administer and therefore more widely accepted are a critical public health need. Here, using a relevant animal model, we have shown that replicative oral AdV vaccine vectors can help make vaccination against major respiratory diseases more available, better accepted and therefore more effective. These results could be of major importance in the coming years in the fight against emerging diseases such as COVID-19.

## Introduction

Human influenza is an acute respiratory disease caused by influenza A and B viruses. Each year, influenza seasonal epidemics are responsible for 3 to 5 million cases of severe illness leading to 300,000 to 500,000 deaths (1). Sporadically, pandemics originating from animal influenza strains occur. In those cases, the absence of pre-existing immunity results in an increased severity and mortality that can be dramatic, as shown by the Spanish flu 1918 pandemic (1). Influenza A and B are enveloped viruses with a genome consisting of eight single-strand negative RNA segments. Two of them encode the viral surface glycoproteins hemagglutinin (HA) and neuraminidase (NA). HA consists of a stalk domain surmounted by a globular head participating to viral entry in host cells (2), while NA contributes to viral spread (3). Both these proteins contain the main epitopes involved in protective antibodies induced by infection or vaccination. They are also antigenically the more variable proteins of the virus. Since influenza viruses are subject to continuous antigenic changes, seasonal influenza vaccines must be adapted every year according to predictions based on circulating strains surveillance (4). The efficacy of these vaccines is hence variable and can be very low, depending on their adequacy with the respect to circulating strains (5). In addition, influenza vaccine production is a time-consuming process prolonging the delay between vaccine strain determination and operative use in the field, which increases the mismatch risk.

Improved influenza vaccination strategies are an important public health challenge. Much research has focused on the development of vaccines giving protection against antigenically distant viruses (6). Among the antigen targets for development of such a universal influenza vaccine, the stalk domain of HA, which is a well conserved part of the protein is promising (7). In parallel, new methods of immunization, such as the mucosal route, could improve vaccine efficacy by inducing immune defences at the portal of virus entry. Furthermore, mucosal administration, and especially the oral route, could reduce costs, be more practical, and improve patient compliance, thereby increasing vaccine coverage (8, 9). However, in the digestive tract, immunity is contained by potent regulatory mechanisms avoiding an inappropriate response against foodstuff or microbiota antigens. To create the inflammatory conditions essential to induce an effective immune response, oral vaccines require therefore efficient delivery systems and powerful adjuvants (9).

Adenovirus (AdV) vectors have features that can fulfill these functions. In particular, replication-competent AdV could provide a potent adjuvant effect through the induction of cytokines and co-stimulatory molecules (10), while producing large amount of antigens. Oral wild type human AdV-4 and -7 based vaccines have been used for decades to protect US military trainees against the severe respiratory diseases caused by these same viruses (11, 12). These vaccines have a good efficacy and safety profile, making them attractive candidates as vectors for oral vaccine platform development. Replication-competent human AdVs have been used as vectors for oral vaccine in several studies, notably against influenza (13–16). However, the results of most of these investigations were rather disappointing. One reason could be that most of these preclinical researches were performed with human AdVs in laboratory animals, i.e., in poor replicative conditions because of the restricted host specific replication of AdVs (17). The use of mouse AdV type 1 (MAV-1) in the mouse, its natural host, permits to study AdVs infection in replicative conditions (18). We recently showed that MAV-1 oral administration in mice reproduces the homologous protection observed in humans with AdV-4 and AdV-7 based oral vaccines (19). In the present study, we exploited this mouse model to investigate the potential of replication-competent AdVs as vectors for oral vaccination against influenza. We used MAV-1 replication-competent recombinants expressing either the full length (FL) or the truncated “headless” (HL) HA (7) from a mouse-adapted viral strain of influenza to evaluate the protection that these oral vaccines conferred against an intranasal influenza infection. We showed that unlike MAV-1 expressing HL HA, which does not generate any detectable adaptive immune response against influenza, the FL HA expressing vector elicits a specific and neutralizing humoral immune response and confers complete protection against clinical signs and lung viral replication from a subsequent respiratory influenza infection.

## Results

### Design and generation of HA expressing MAV-1 recombinants

We constructed two MAV-1 recombinant vectors expressing either the FL HA of influenza A PR8 or a truncated form of this protein (HL HA), in which the globular head was deleted while the stalk domain was kept unchanged (Fig. 1A). Using a bacterial artificial chromosome (BAC) containing the entire MAV-1 genome (20), the FL or HL HA ORFs were inserted downstream of the protein IX (pIX) sequence. Since AdVs cannot accommodate large transgenes, the pIX stop codon was replaced by a furin 2A cleavage sequence linking HA transgene expression to transcription of pIX without addition of transcriptional control regions (21) (Fig. 1A). The resulting mRNA allowed to give rise to distinct HA and pIX proteins through enzymatic cleavage of the furin 2A site. The constructions of HL and FL HA MAV-1 recombinants were checked by ApaI or SspI enzymatic restrictions of the BAC genomes, showing the expected profiles (Fig. 1B), and confirmed by sequencing of recombination regions (data not shown).

**FIG 1.**
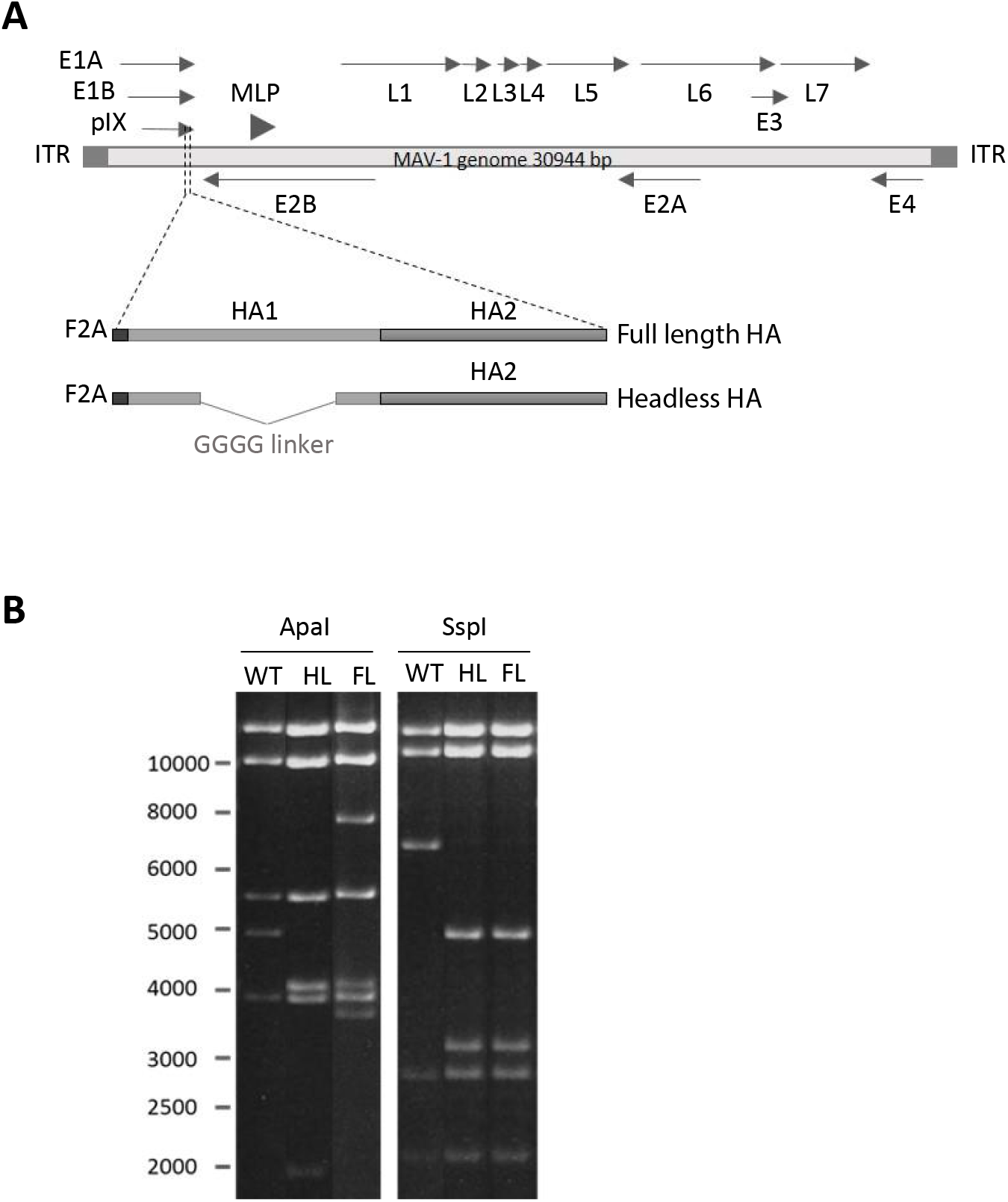
Construction and characterization of MAV-1 recombinant expressing either the full length (FL) or the headless (HL) hemagglutinin (HA) of influenza PR8. (A) Schematic representation of the strategy used to produce the recombinant MAV-1 strains. A furin 2-A (F2A)-FL or HL HA cassette was inserted instead of the stop codon of the pIX ORF by homologous recombination using a bacterial artificial chromosome (BAC) containing the complete MAV-1 genome. (B) Verification of the molecular structure. Wild type (WT), HL HA or FL HA MAV-1 BAC were purified from bacteria and digested with *Apa*I or *Ssp*I digestion. The restriction fragments were then separated by electrophoretic migration and revealed by ethidium bromide and UV exposure. Marker sizes in Kbp are indicated on the left.

These recombinant MAV-1 had a growth deficit in 3T6 cells compared to the wild type (WT) MAV-1 (data not shown), probably due to the increased genome size. Expression of both the HL and FL HA was checked by immunofluorescence on infected cells. We observed a clear staining of MAV-1 FL HA infected cells, but a weaker signal from MAV-1 HL HA infected cells (Fig. 2A). This observation was not surprising as we used a primary polyclonal serum from mouse immunized with inactivated influenza, which is expected to mainly contain antibodies against the immunodominant globular head of HA rather than against the stalk domain. We further confirmed the expression of HL HA by the MAV-1 recombinant by western blotting (Fig. 2B).

**FIG 2.**
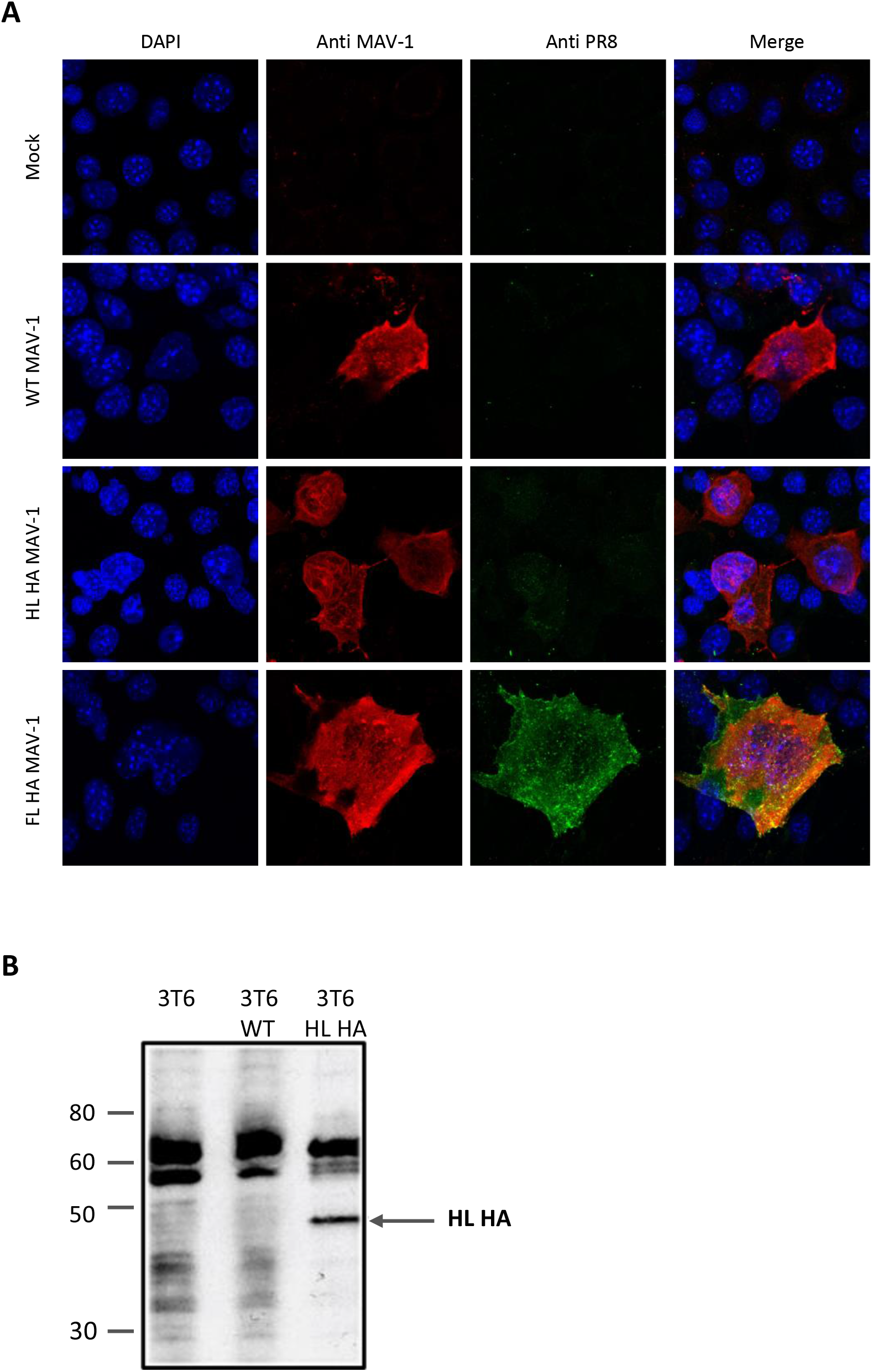
Assessment of FL HA and HL HA expression by MAV-1 recombinants. (A) WT, FL HA or HL HA MAV-1 were grown in 3T6 cells until first cytopathic effects were seen (approximately 5 days post-infection). Infected cells were then fixed and labelled using a primary rabbit antiserum to MAV-1 structural proteins (57) and a mouse polyclonal antiserum to PR8 structural proteins, followed by conjugated secondary antibodies. Nuclei were stained with DAPI. (B) 3T6 cells were uninfected or infected with WT or HL HA MAV-1. When cytopathic effects appeared, cells were collected, and proteins were submitted to western blotting as described in the methods. HL HA was detected with a mouse polyclonal antiserum to PR8 structural proteins as described in the methods. Positions of molecular weight standards in kDa are indicated on the left.

### Oral immunization with MAV-1 FL HA protects against a lethal respiratory challenge with influenza

To evaluate the ability of orally administered AdV vectored vaccines to confer a protective immune response against influenza infection, mice were immunized by gavage with 104 TCID50 of MAV-1 HL HA or MAV-1 FL HA recombinants or, as negative controls, the same dose of MAV-1 WT or PBS only (mock) (Fig. 3A). Mice were observed and weighed every other day from the day before immunization until day 9 post-immunization. None of the mice developed any clinical sign consecutive to vaccination (data not shown). At day 27 post-immunization, serum was collected, and PR8 specific (Fig. 3B) and neutralizing (Fig. 3C) antibodies were measured by in-cell ELISA and seroneutralization assay respectively. In the MAV-1 HL HA group, similar to mock and MAV-1 WT control groups, no PR8 specific or neutralizing antibodies were detected. In contrast, we detected significant levels of specific and neutralizing antibodies in the MAV-1 FL HA group, showing that administration of this recombinant generates a humoral immune response against influenza.

**FIG 3.**
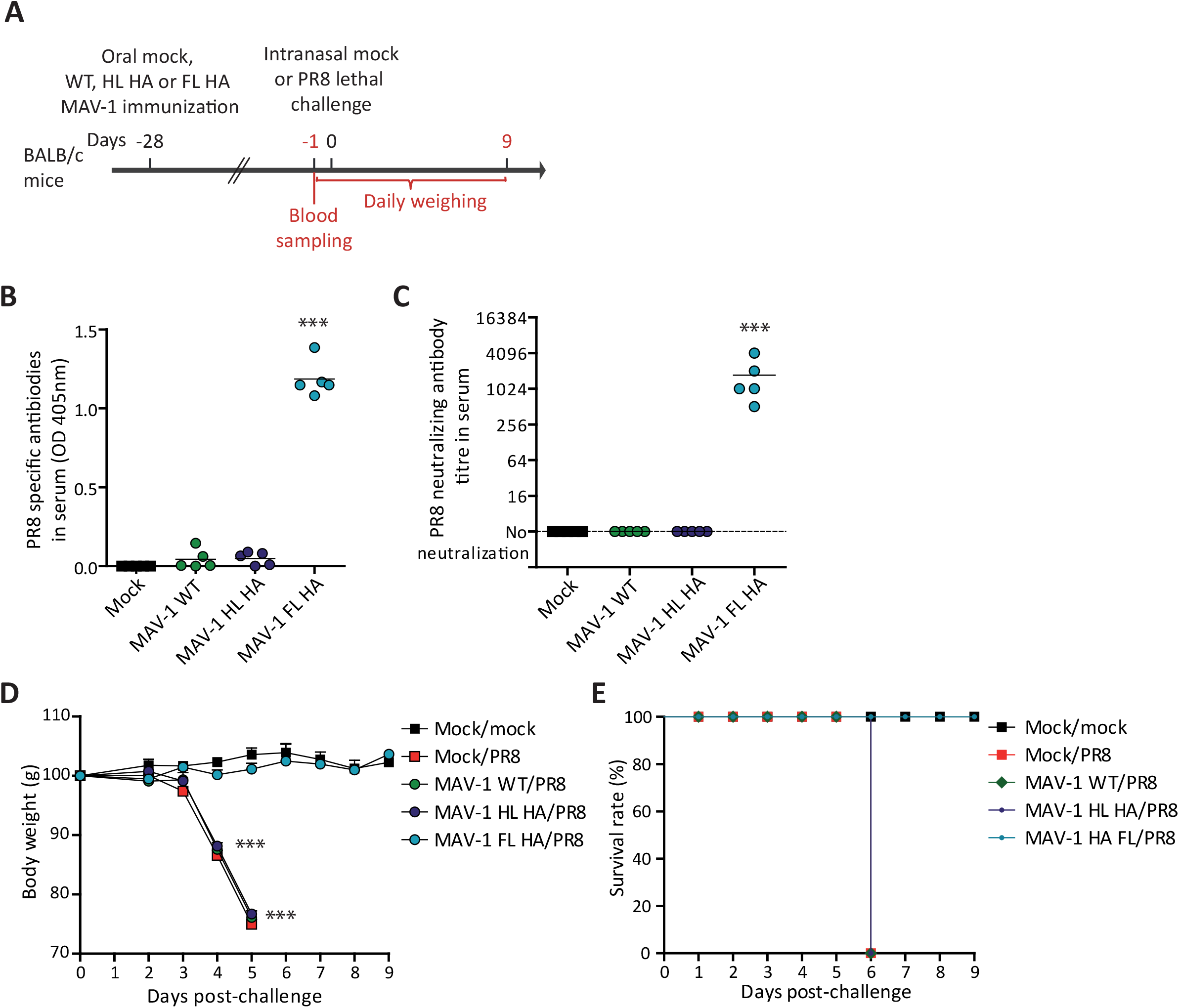
Evaluation of the protection induced by HL HA MAV-1 and FL HA MAV-1 oral immunization against a respiratory influenza challenge. (A) Eight-week-old female BALB/c mice were orally immunized with 104 TCID50 of WT, HL HA or FL HA MAV-1 or with PBS as control (mock). At day 28 post-immunization, mice were challenged or not by intranasal administration of 2.5×103 PFU of influenza PR8. Blood was collected the day before challenge. Mice were weighed and observed daily from the day before challenge up to 9 days post-challenge (n=5 in each group). (B-C) PR8-specific (B) and PR8-neutralizing antibody titres (C) at day 27 post-immunization. OD, optical density. (D) Weight of mice after influenza virus challenge. For each time point, means were compared to the mean of the mock/mock group. (E) Survival curve until day 9 post-challenge. The data presented are either the means for 5 mice ± SEM (D) or individual data and means (B and C). *P* values are relative to comparison with the mock/mock group. ***, *P*<0.001.

To know if this antibody response correlates with clinical protection, BALB/c mice were intranasally challenged or not with a lethal dose (2.5 103 PFUs) of the mouse adapted influenza PR8 strain (Fig. 3A). They were then monitored by daily observation and weighing from the day before challenge up to 9 days post-challenge. From day 4 post-challenge, non-immunized challenged mice (mock/PR8), MAV-1 vector immunized challenged mice (MAV-1 WT/PR8), and MAV-1 HL HA immunized challenged mice (MAV-1 HL HA/PR8) developed clinical signs characterized by weight loss (Fig. 3D), ruffled fur, hunched posture, dehydration, dyspnea and lethargy. At day 5 post-challenge, all these mice died from the disease or were euthanized for ethical reasons (> 20% weight loss). In contrast, mice that were previously orally immunized with the MAV-1 FL HA recombinant (MAV-1 FL HA/PR8) did not display any weight loss (Fig. 3D) or clinical sign after influenza challenge, similar to the non-immunized non-challenged group (mock/mock). Together, these results show that a single oral immunization with a replication-competent AdV vector expressing the entire HA of influenza elicited a humoral immune response and conferred full protection against clinical signs due to a subsequent respiratory influenza PR8 lethal challenge. In contrast, the same MAV-1 vector expressing the truncated HL HA failed to confer any detectable humoral response or clinical protection against this lethal PR8 challenge.

### Protection conferred by oral immunization with MAV-1 FL HA blocks innate immune inflammation and does not rely on specific T cells

To further investigate the protection conferred by oral administration of MAV-1 vectors, mice were mock-immunized or immunized with 104 TCID50 of MAV-1 HL HA, MAV-1 FL HA or MAV-1 WT before being mock or influenza challenged with a sublethal dose of PR8 (25 PFU), in order to analyse inflammation and viral replication in the lungs one week post-challenge (Fig. 4-6).

**FIG 4.**
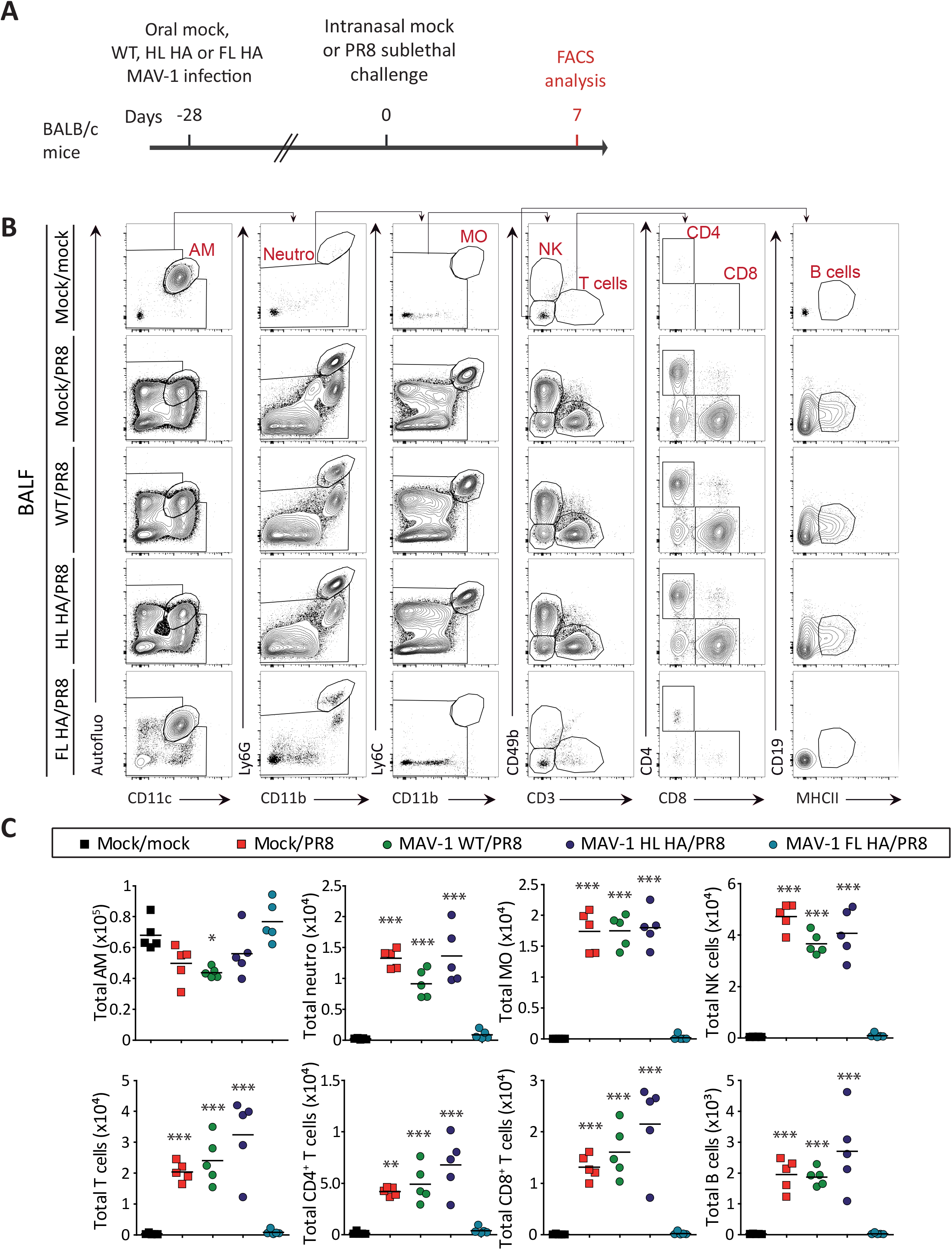
Characterization of immune cell populations in bronchoalveolar lavage after HL HA MAV-1 and FL HA MAV-1 oral immunization and influenza PR8 respiratory challenge. (A) Eight-week-old female BALB/c mice were orally immunized with 104 TCID50 of WT, HL HA or FL HA MAV-1 or with PBS as a control (mock). At 28 days post-immunization, mice were challenged or not by intranasal administration of 25 PFU of PR8 influenza. Mice were euthanized at day 7 after intranasal challenge. (B) Gating strategy and (C) total numbers of immune cell populations in bronchoalveolar lavages. *P* values are for comparisons between all pairs of groups. *, *P*=0.05; **, *P*=0.01; ***, *P*=0.001. AM, alveolar macrophages; Neutro, neutrophils; MO, monocytes.

First, we characterized the immune cell infiltration in the bronchoalveolar lavage fluids (BALF) (Fig. 4) and in the lung parenchyma (Fig. 5) at day 7 post-challenge. In mock/PR8, MAV-1 WT/PR8 and MAV-1 HL HA /PR8 groups, we observed a massive cellular infiltrate primarily made of NK cells, neutrophils and inflammatory monocytes (Ly6Chi monocytes) in the BALF (Fig. 4B and C). In the lung parenchyma of mice from the same groups, we observed an increased number of alveolar macrophages (AM) and a massive infiltrate of inflammatory monocytes and natural killer (NK) cells (Fig. 5 B and C). This infiltration of innate immune cells was completely abolished in the MAV-1 FL HA/PR8 group. Moreover, in the MAV-1 FL HA/PR8 mice, the total numbers of B cells, CD4+ T cells and CD8+ T cells were comparable to the mock/mock group, suggesting that the protection observed against influenza essentially relies on systemic humoral immune response. In contrast, in the airways of mock/PR8, MAV-1 WT /PR8 and MAV-1 HL HA/PR8 mice, we observed significant amount of B cells, CD4+ T cells and CD8+ T cells (Fig. 4B and C), and in the lung tissue of MAV-1 WT/PR8 and MAV-1 HL HA/PR8 groups, significant levels of CD8+ T cells (Fig. 5B and C) displaying mainly an effector memory (EM) phenotype (Fig. 6B and C). In parallel, we estimated the amount of HA stalk specific CD8+ T cells in lung parenchyma by MHC I tetramer labelling. This analysis showed higher numbers of HA specific cytotoxic T cells in MAV-1 WT/PR8 and in MAV-1 HL HA/PR8 groups than in MAV-1 FL HA/PR8 group that was however fully protected (Fig. 6B and C). These results indicate that the MAV-1 FL HA vaccine completely protected against lung inflammation consecutive to influenza respiratory challenge. This protection against influenza seems to not rely on lung cellular innate immunity or on specific CD8 T cells. On the contrary, oral administration of the MAV-1 HL HA vector did not protect against lung inflammation caused by an influenza challenge.

**FIG 5.**
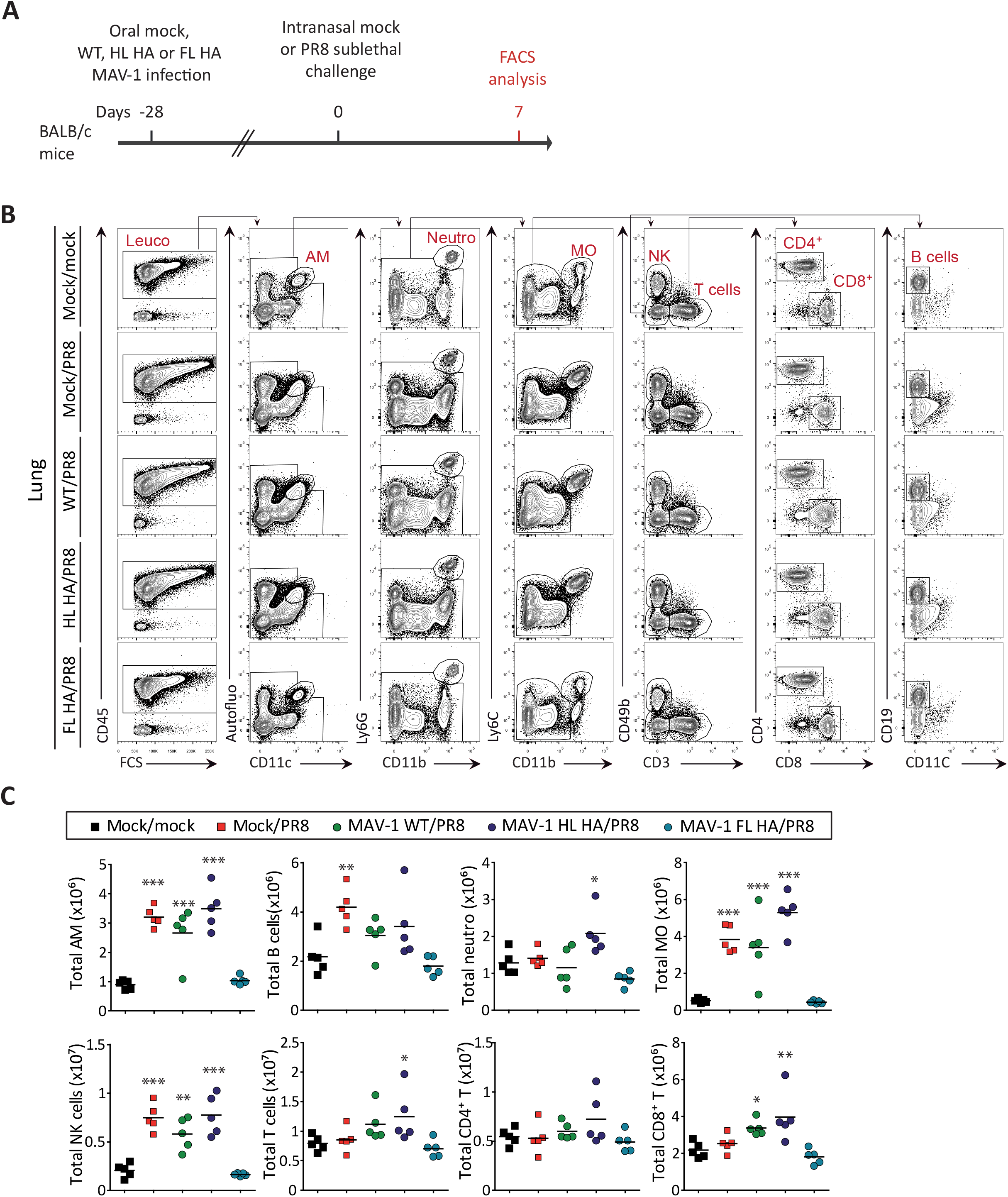
Characterization of lung parenchyma immune cell populations after HL HA MAV-1 and FL HA MAV-1 oral immunization and influenza PR8 respiratory challenge. (A) Eight-week-old female BALB/c mice were orally immunized with 104 TCID50 of WT, HL HA or FL HA MAV-1 or with PBS as a control (mock). At 28 days post-immunization, mice were challenged or not by intranasal administration of 25 PFU of PR8 influenza. Mice were euthanized at day 7 after intranasal challenge. (B) Representative flow cytometry dot plots (B) and total numbers (C) of immune cell populations in lung parenchyma. *P* values are for comparisons between all pairs of groups. *, *P*<0.05; **, *P*<0.01; ***, *P*<0.001 (one-way ANOVA and Bonferroni post-tests). AM, alveolar macrophages; Neutro, neutrophils, MO, monocytes; Leu, leukocytes.

**FIG 6.**
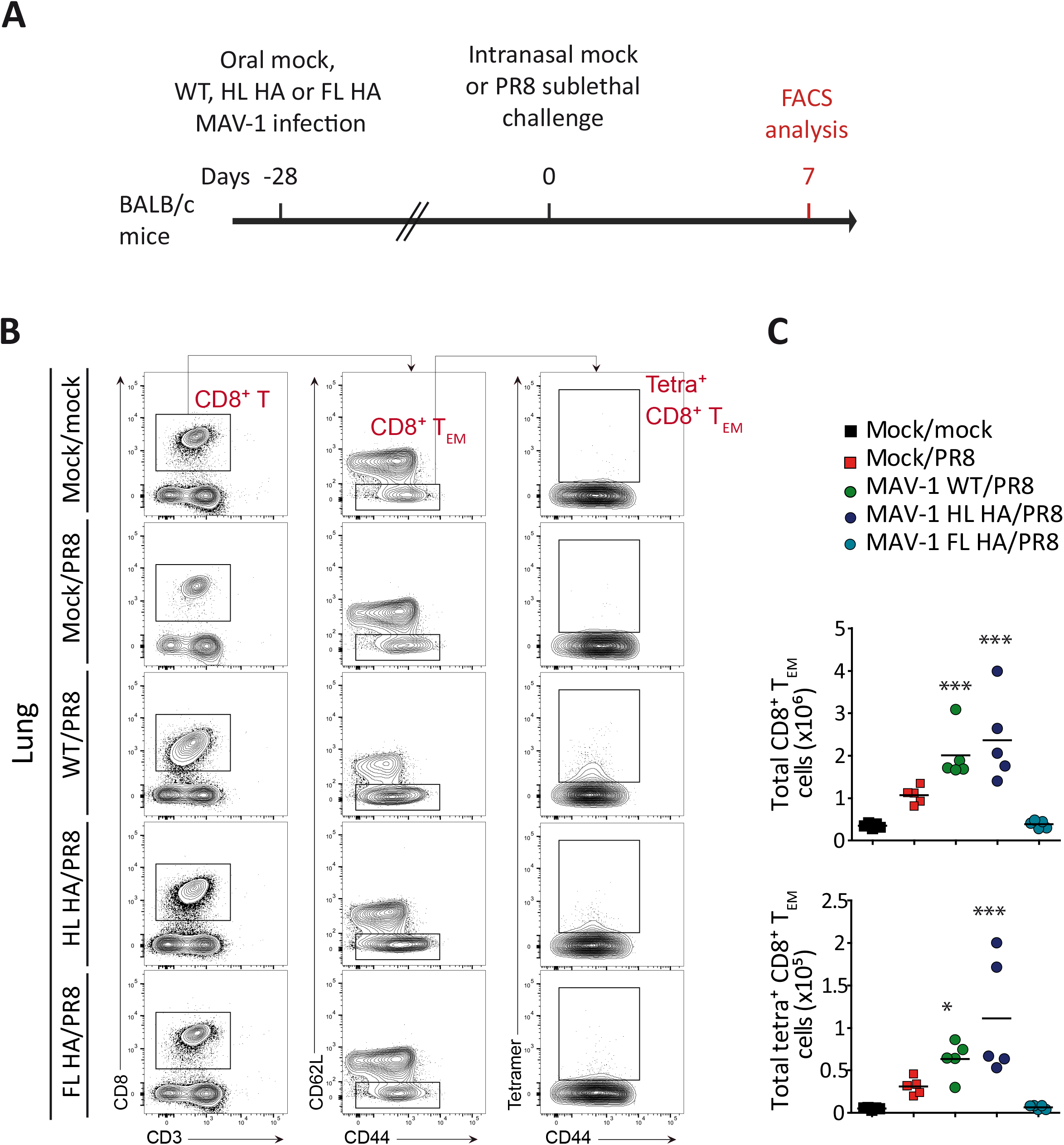
Quantification of lung parenchyma influenza specific CD8+T cells. (A) Eight-week-old female BALB/c mice were orally immunized with 104 TCID50 of WT, HL HA or FL HA MAV-1 or with PBS as a control (mock). At 28 days post-immunization, mice were challenged or not by intranasal administration of 25 PFU of PR8 influenza. Mice were euthanized at day 7 after intranasal challenge. Representative flow cytometry dot plots (B) and total numbers (C) CD8+T cells and of effector memory influenza specific CD8+T cells in lung (cells were pre-gated as live CD45+ non-autofluorescent CD11c-CD3+CD8+CD62L-CD44hi). *P* values are for comparisons between all pairs of groups. *, *P*<0.05; **, *P*<0.01; ***, *P*<0.001 (one-way ANOVA and Bonferroni post-tests). TEM cells, effector memory T cells; Tetra, Influenza A HA tetramer positive cells.

Finally, we evaluated the protection against influenza replication in the lung. For this purpose, we titrated the virus in the lung tissue by plaque assay one week post-sublethal challenge (Fig. 7A). We detected high amounts of influenza viruses in the mock/PR8, MAV-1 WT/PR8 and MAV-1 HL HA/PR8 groups, though the levels tended to be lower in this last group. In contrast, in the MAV-1 FL HA /PR8 group, no PR8 virus was detected (Fig. 7B). These results show that, while the MAV-1 HL HA fails to prevent influenza replication in the lung, the MAV-1 FL HA vaccine fully protects against respiratory infection and viral replication.

**FIG 7.**
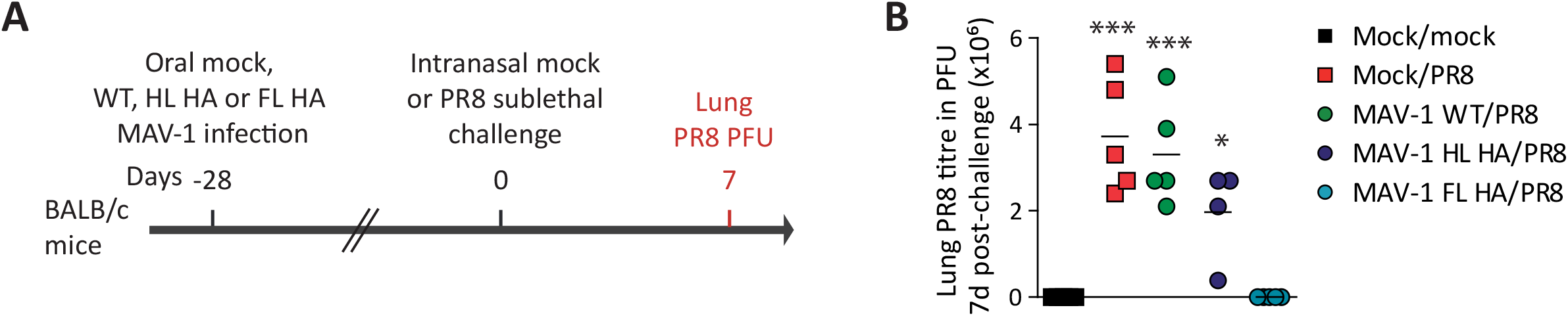
Quantification of influenza PR8 titres in lungs after HL HA MAV-1 and FL HA MAV-1 oral immunization and influenza PR8 respiratory challenge. (A) Eight-week-old female BALB/c mice were orally immunized with 104 TCID50 of WT, HL HA or FL HA MAV-1 or with PBS as a control (mock). At 28 days post-immunization, mice were challenged or not by intranasal administration of 25 PFU of PR8 influenza. Mice were euthanized at day 7 after intranasal challenge. (B) Numbers of PR8 PFU in the lungs at day 7 post-challenge. *P* values are for comparisons between all pairs of groups. *, P<0.05; ***, P<0.001 (one-way ANOVA and Bonferroni post-tests).

## Discussion

The development of influenza vaccines is a major public health issue (22). While the identification of conserved antigens is a priority in order to improve the efficacy of vaccines in the face of seasonal variations of the virus (23, 24), the development of new routes of administration seems equally important in order to broaden the adherence of target populations and facilitate the organisation of vaccination campaigns (4, 25). Indeed, in light of the COVID-19 crisis, it appears that economic and logistical considerations can crucially impact the deployment of vaccination at the worldwide scale (26).

Here we established a mouse model to study and evaluate replication-competent AdV-vectors as oral vaccine platforms for influenza, and potentially for other infectious respiratory diseases such as the one caused by SARS-CoV-2. To this end, we constructed recombinant MAV-1 expressing either the FL or the HL HA of the mouse adapted strain of influenza PR8. Our results showed that MAV-1 replication competent AdV vector expressing the FL HA was able to elicit a potent specific and neutralizing systemic antibody response associated with full protection against a lethal influenza challenge. In contrast, the same vector expressing HL HA, which is described as a “universal” influenza antigen (7, 27) failed to generate any detectable protection or immune response against influenza. Oral administration of MAV-1 FL HA vaccine completely prevented mortality, weight loss or other visible signs of disease, and inhibited lung viral replication. Our results suggest that the protection observed mainly relies on systemic antibodies which would transude through the respiratory mucosa and block virus entry so efficiently that no other immune response is required. In accordance with these results, the absence of viral replication detected by plaque assay on lung tissue one week after the challenge suggests sterile immunity. Although earlier time-points should be tested to confirm these results, that level of immunity would constitute a substantial benefit, since it could allow breaking of the transmission chain. It is notable that an oral virus vectored vaccine triggers so strong a systemic humoral immune response, because oral administration is first expected to induce local mucosal immunity (28). The level of serum antibodies is considered as the gold standard correlate of protection for influenza (29–31) and many other diseases (32, 33). The ability to induce such high levels of specific antibody in serum is thus promising for further vaccine evaluation and development, as well as for epidemic or pandemic management (34).

In contrast, the HL HA vaccine did not generate any significant immune protection. Considering that most epitopes involved in the protective neutralizing antibody response are contained in the globular head (35), a less efficient immune response was expected with this antigen (23, 36). A previous study using HL HA in a vaccine regimen based on two injections of DNA plasmid vector, followed by one injection of Freund adjuvanted VLP, showed that this antigen was able to give rise to specific and neutralizing antibodies cross reacting against different influenza subtypes (7). However, that humoral response was only observed in some recipient mice, while others did not respond. We postulated here that a replication competent AdV oral vaccine vector could potentially improve the efficiency of the HL HA based vaccination strategy by allowing *in vivo* amplification of the antigenic transgene and stimulating mucosal immunity (37). However, these theoretical advantages were refuted by our results.

Surprisingly, after influenza virus challenge, total CD8 T cells and HA stalk domain specific CD8 T cells were significantly higher in the WT MAV-1 and HL HA MAV-1 groups than in the mock group (Fig. 5C and 6C). This suggests that the vector itself contributed to the development of a specific cytotoxic response and illustrates the potential ability of AdV vectors to enhance immunity against a heterologous challenge and to orientate it toward CD8 T cell response. This better CD8 response also seemed to correlate with lower lung viral load (Fig. 7). A similar boost of heterologous immunity was recently made in the context of other viral infections by our group (38). Considering these observations, development of vectored-vaccines based on replication competent AdVs could also be advantageous for an improved T cell immune response. Cellular immunity to influenza raises growing interest in the field of universal influenza vaccine researches (24, 39–41). T antigens endure less immune pressure than B antigens and should be more conserved for different influenza strains. Several studies suggest that cytotoxic response could provide at least partial protection (42, 43). Since our HL HA vaccine was not efficient, other antigens known to trigger a good cytotoxic response could be considered. For example, influenza nucleoprotein specific CD8+ T cells partially prevent influenza mortality in mice (40).

Finally, recently AdV vectors have emerged as powerful weapons to oppose to the SARS-CoV-2 pandemic. The worldwide use of these platforms to control COVID-19 confirms and supports the feasibility of AdVs for immunization against other infectious diseases, particularly influenza. These vaccine platforms are indeed efficient and easy to adapt and produce. Moreover, as compared to mRNA technologies, they are cheaper, and their conservation does not necessitate extremely low temperatures. However, AdV based vaccines remain optimisable. In that regard, the use of replication competent vectors through the oral route could offer critical improvement in production, distribution, and administration, thus contributing to vaccine access and equity. Furthermore, needle free administration would certainly help to overcome vaccine hesitancies. Finally, from an immunological point of view, the potential to confer sterile immunity would allow attaining herd immunity faster.

In this study, we showed in a mouse model that an oral replicative AdV based vaccine provided complete protection against a lethal respiratory challenge without generating any detectable adverse effect. Although these encouraging results were obtained with a highly immunogenic antigen and a perfectly homologous challenge, they encourage the study of replicating AdVs as vectors for the development of oral vaccines against influenza or other respiratory diseases, including COVID-19. Several studies have investigated the potential of replication-competent AdV oral vaccine vectors, notably against influenza (44–46). However, those researches generally used human AdV in animals, i.e., in non-replicative conditions, which limits applicability of results in terms of animal to human translation. Phase I studies have also been done in humans, but oral AdVs were only used in combination with intramuscular inactivated seasonal vaccine boost (47). To our knowledge, an animal model allowing study of replication competent AdV vaccine vectors in a permissive host, such as MAV-1 in mouse, has been studied to date (19, 48). This work establishes an original and useful tool to better evaluate AdV vaccine platforms *in vivo*. This opens new possibilities for preclinical evaluation of replication competent AdV vectors in terms of safety and efficacy. Reliable animal models are needed to provide sound data on the risks and benefits associated with new generation vaccine platforms, so that these technologies can be used in an informed manner when an emergency requires rapid responses.

## Materials and Methods

### Animals

All animal work complied with relevant European, federal, and institutional policies. The Committee on the Ethics of Animal Experiments of the University of Liège reviewed and approved the protocol (permit number 1526). Female BALB/c mice were purchased from Envigo. All mice were housed in the University of Liège, Department of Infectious Diseases. The animals were infected orally or intranasally with MAV-1 or PR8 influenza, when 7-8 and 11-13 weeks old, respectively. For oral infections, 104 TCID50 of MAV-1 diluted in 250 µL of phosphate-buffered saline (PBS) were administered into the oesophagus with a steel feeding gavage needle. For intranasal infections, 25 (sub-lethal dose) or 2.5×103 (lethal dose) PFU of PR8 influenza diluted in 50 µL of PBS were inoculated under mild isoflurane anaesthesia. To evaluate clinical signs of infection, we monitored and weighed mice daily for 10 days. Ruffled coat, loss of activity and weight loss below 20% were considered as signs of moderate pain and distress; hunched posture, dyspnea, lethargy and weight loss greater than 20% were considered as signs of severe pain and distress and constituted endpoints criteria (49).

### Virus and cells

We used the pmE101 wild type strain of MAV-1 (50, 51) and the mouse adapted strain of influenza A/Puerto Rico/8/1934 (H1N1, PR8) (52). MAV-1 and PR8 were grown on mouse 3T6 cells and Madin-Darby Canine Kidney (MDCK) cells respectively, cultivated in Dulbecco’s modified Eagle’s medium (DMEM; Life Technologies), supplemented with 2 mM glutamine, 100 U penicillin mL-1, 100 mg streptomycin mL-1 and 5% heat-inactivated fetal calf serum (FCS) at 37°C in an atmosphere of 5% CO2. For PR8 growth, TPCK-treated trypsin (2 mg/ml) was added to culture medium.

### Construction of HA expressing MAV-1 recombinant vectors

FL HA was amplified with HA_fwd 5’-GACCTTCTCAAGTTGGCAGGAGACGTTGAGTCCAACCCTGGGCCCATGAAGGCAAACCTACTGGTCCTGTTAAG-3’ and HA_rev 5’-TTTTTATTAAACATAAAGCGCGTGAGCATGCATCTTTATTTGGGATCAGATGCATATTCTG CACTG-3’ primers by RT-PCR from influenza PR8 genome. HL HA was amplified from FL HA by 5 successive rounds of PCR using HA_rev as reverse primer and the 5 successive forward primers: HL_HA1 fwd 5’-GGTGGTGGTGGTTGTAACACGAAGTGTCAAACA-3’; HL_HA2 fwd 5’-TGTGACAGTGACACACTCTGTTAACCTGCTCGAAGACAGCCACAACGGAAAACTATGTGGTGGTGGTGGTTGTAACAC-3’; HL_HA3 fwd 5’-GTATAGGCTACCATGCGAACAATTCAACCGACACTGTTGACACAGTACTCGAGAAGAATGTGACAGTGACACACTCTG-3’; HL_HA4 fwd 5’-AAACCTACTGGTCCTGTTAAGTGCACTTGCAGCTGCAGATGCAGACACAATATGTATAGGCTACCATGCGAAC -3’; HL_HA5 fwd 5’-<u>CAAACTTTGAATTTTGACCTTCTTAAGCTTGCGGGAGACGTCGAGTCCAACCCTGGGCCC</u> ATGAAGGCAAACCTACTGGTCCTGTTAAG -3’. These cassettes were cloned downstream and in frame with the MAV-1 pIX ORF. A sequence coding for a furin 2A cleavage site (underlined in the HL_HA5 fwd primer) was added between pIX and HA sequences to allow release of HA proteins. Recombinants were produced using *Escherichia coli* SW102 containing the pKBS2.MAVLJ1 wild type (WT) bacmid coding for the entire MAV-1 WT (53) and prokaryotic recombination technologies as described before (54). Briefly, a GalK cassette, amplified with GalK fwd 5’-GACCTTCTCAAGTTGGCAGGAGACGTTGAGTCCAACCCTGGGCCCCCTGTTGACAATTAATCATCGGCA-3’ and GalK rev 5’-TTTTTATTAAACATAAAGCGCGTGAGCATGCATCTTTATTTGGGACTCAGCACTGTCCTGC TCCTT-3’, was introduced in placethe of pIX stop codon to allow positive selection of recombinants on a minimal medium, then this insert was replaced by the FL HA and HL HA cassettes allowing counter-selection on a DOG medium toxic for non-recombinants. The screening of recombinants was made by PCR and confirmed by restriction profile. After BAC purification, the mutated MAV-1 genome was excised from the plasmid and transfected in 3T6 cells in order to rescue the recombinant viruses.

### Restriction profile

100 µl of an overnight culture of SW102 *E. coli* containing MAV-1 WT or recombinant BAC were diluted in 10 ml Luria–Bertani (LB) medium supplemented with chloramphenicol (12.5 mg/ml) at 32°C in a shaking incubator. 1 µg DNA of BAC miniprep from this culture was digested with ApaI or SspI for 2h at 25 and 37°C respectively. The digestion products were separated by electrophoresis on a 0.8% agarose gel.

### Production of PR8 antiserum

To produce anti-PR8 serum, the virus (108 PFU/mL) was incubated for 72 h in 0.02% formaldehyde at 37°C. Formaldehyde was then removed by dialysis in PBS (20 KDa dialysis cassette, Thermo Scientific). Mice were immunized by intraperitoneal administration of 150 µL of this inactivated viral stock. One month later, mice were euthanized, serum was collected and stored at -20°C.

### Western blotting

3T6 cells plated in 12 wells were either mock-infected or infected with WT MAV-1 or MAV-1 HL HA. When cytopathic effects appeared, cells were washed with PBS, then scraped, collected and centrifuged at 1000 *g* for 20 min at 4°C. The pellet was suspended in PBS, then mixed with 4X Laemmli sample buffer supplemented with 10% 2-betamercaptoethanol. The cell lysates were incubated 5 min at 95°C. Proteins were separated by SDS-PAGE in a Mini-PROTEAN TGX gel 4-15%, then transferred on polyvinyl difluoride membrane. Membranes were blocked for 2 h at room temperature in PBS containing 0.1% Tween 20 and 5% dry milk and incubated overnight at 4°C with anti-PR8 polyclonal mouse serum; they were then washed three time in PBS containing 0.1% tween 20 and incubated 1h with a rabbit anti-mouse antibody conjugated with horseradish peroxidase (HRP) (Dako). The activity of HRP was revealed by enhanced chemiluminescence (Perkin-Elmer).

### Indirect immunofluorescence staining of adherent cells

3T6 cells plated on coverslip were either mock-infected or infected with MAV-1 WT, MAV-1 HL HA or MAV-1 FL HA. When cytopathic effect was observed (approximately 5 days p.i.), cells were washed once in PBS, fixed in 4% paraformaldehyde for 10 min at room temperature, washed 2 times in PBS and blocked for 30 min with PBS, 10% FCS, 300 mM glycine. Cells were then incubated overnight at 4°C with a primary rabbit antiserum to MAV-1 structural proteins (55) diluted 2000-fold in the blocking solution and a mouse polyclonal antiserum to PR8 structural proteins diluted 500-fold. Cells were washed 3 times in PBS and incubated at room temperature for 45 min with a goat anti-mouse Ig polyclonal secondary antibody conjugated with Alexa-fluor 488 and a goat anti-rabbit Ig polyclonal secondary antibody conjugated with Alexa-fluor 568. After 3 washes with PBS, nuclei were stained with 4’-6-diamidino-2-phenylindole (DAPI), then washed three times in PBS. Coverslips were mounted in ProLong Glass Antifade Mountant (ThermoFisher) on glass plates. Z-stack images were taken using the Leica SP5 confocal microscope and compiled using ImageJ Software (56).

### Quantification of PR8 specific antibodies by in-cell ELISA

3T6 cells were plated in 96-well plates at 5×103 cells per well, infected with 500 TCID50 of MAV-1 per well, and incubated at 37°C, 5% CO2. When cytopathic effect was observed (approximately 5 days p.i.), cells were fixed in 2% paraformaldehyde for 20 min at RT, washed 2 times in PBS and blocked for 30 min with PBS, 10% FCS, 300 mM glycine. Cells were then incubated at 4°C for 2 h with mouse sera diluted 100-fold in the blocking solution. Cells were washed 3 times in PBS and incubated at room temperature for 30 min with a goat anti-mouse Ig polyclonal secondary antibody conjugated with alkaline phosphatase (0.5 µg/mL, Sigma). After 3 washes with PBS, p-nitrophenyl phosphate (Sigma) was added and incubated for one hour, then the reaction was stopped with 1M sodium hydroxide. Absorbance was measured at 405 nm using a benchmark ELISA plate reader (Thermo). Signals detected from mock infected cells were used as background and this background was subtracted from the corresponding values obtained with MAV-1-infected wells.

### In vivo samples collection and processing

Mice were euthanized by isoflurane inhalation. After euthanasia, the trachea was catheterized and BAL was performed by 2 consecutive flushes of the lungs with 1 mL ice-cold PBS containing Complete Protease Inhibitor Cocktail (Roche). Cell density in BAL fluid (BALF) was evaluated using a hemocytometer after Tuerk solution staining (Sigma-Aldrich). Differential cell counts in BALF were determined by flow cytometry analysis. For lung tissue and cells suspension preparations before plaque assay and FACS analysis, lungs were perfused with ice-cold PBS through the right ventricle. Then, lung lobes were collected into a C-Tube (Miltenyi) containing complete HBSS medium, 1 mg/mL collagenase D (Roche) and 50 µg/mL DNase I (Roche), processed with a gentleMACS dissociator (Miltenyi). One third of this homogenate was kept at -80°C to be titrated latter; the remaining two-thirds were incubated for 30 min at 37°C, then washed, treated to lyse of the erythrocytes (Red Lysis Buffer, eBioscience), and strained through a 70-µm filter. Cell density was evaluated using a hemocytometer after Tuerk solution staining (Sigma-Aldrich). Differential cell counts in lung parenchyma were determined by flow cytometry analysis.

### Flow cytometry

Labelling of single cell suspensions was performed in FACS buffer (PBS containing 2 mM EDTA, 0.5% BSA and 0.1% NaN3) at 4°C. Cells were first incubated for 20 min with a purified rat IgG2a anti-mouse CD16/CD32 antibody (1/500) to block Fc binding. FACS staining was performed by using different panels of the following fluorochrome-conjugated antibodies against CD45 (30-F11, PE-Cy7, 1/2000), MHC-II (M5/114.15.2, PE-Cy7, 1/2000), CD3e (145-2C11, APC, 1/400), CD8 (53-6.7, PE, 1/1000), CD4 (RM 4-5, V450, 1/500), CD19 (MB19-1, APC-Cy7, 1/500), CD49b (DX5, FITC, 1/500), Ly6G (1A8, APC-Cy7,1/500), CD11c (N418, Alexafluor700, 1/500), CD11b (M1/70, BV711, 1/1500), Ly6C (HK1.4, BV785, 1/1500), CD62L (MEL-14, eFluor 450, 1/1000), CD44 (IM7, PE-Cy7, 1/1000). All antibodies were purchased from eBioscience, BD Biosciences or Biolegend. Dead cells were stained using Fixable Viability Stain 510 (BD Bioscience) or Fixable Viability Dye eFluor 780 (eBioscience). HA A influenza specific CD8+ T cells were stained with H-2K(d) Influenza A HA 533-541 IYSTVASSL tetramer (APC 1/500) obtained from NIH Tetramer Core Facility.

Samples were analysed on a BD LSR Fortessa X-20 flow cytometer equipped with 50-mW violet 405 nm, 50-mW blue 488 nm, 50-mW yellow-green 561 nm, and 40-mW red 633 nm lasers and a ND1.0 filter in front of the FSC photodiode, using BD FACSDiva software. Data were analysed with FlowJo software (Tree Star).

### Determination of influenza PR8 titre from lung tissue

MDCK cells were plated in 6-well wells at 9 × 105 cells per well. When adherent, they were washed three times with PBS then infected with 10-fold serial dilutions of lung homogenates in 1 mL DMEM supplemented with 2 mM glutamine, 100 U penicillin/mL, 100 mg streptomycin/mL and TPCK-treated trypsin (2 mg/ml), then incubated at 37°C, 5 % CO2. After 2 hours, 2 mL of carboxymehylcellulose medium viscosity (0.6 % final concentration) – DMEM was added to the cells. When cytopathic effects were observed (approximately 1-3 days p.i.), cells were fixed in 2% paraformaldehyde for 20 min at RT, washed in PBS and stained with trypan blue.

### Statistical analysis

Data were analysed with GraphPad Prism 5.0 software. For all the experiments, n=5 in each group. For weight curves, the data were analysed by 2-way ANOVA and Bonferroni post-tests. For serological and PR8 titer analysis we used the Wilcoxon–Mann–Whitney rank-sum test. For lung and BALF cellular populations, data were analysed by 1-way ANOVA and Bonferroni post-tests.

## Acknowledgements

E.G. was a research fellow of the “Fund for Research Training in Industry and Agriculture” (FRIA). The authors thank Katherine R. Spindler for helpful advice on the experiments and critical reading of the manuscript. The authors thank Mamadou Diallo for help with confocal analyses. The authors thank the technician and administrative team of the lab for very helpful assistance. The authors thank Michel Bisteau, Eric Destexhe and Philippe Hermand for their support in the design of the experiments and data analysis. The authors thank the NIH Tetramer Core Facility for providing tetramers.

## Contributions

L.G., E.G. and B.M. designed the experiments with the help of S.H.; E.G. performed the experiments and compiled the data; E.G., B.M., and L.G. analysed the data; E.G., B.M. and L.G. prepared the figures; E.G. and L.G. wrote the manuscript; all authors critically revised the manuscript for important intellectual content. All authors had full access to the data and approved the manuscript before it was submitted by the corresponding author.

